# Electrical synapses between inhibitory neurons shape the responses of principal neurons to transient inputs in the thalamus

**DOI:** 10.1101/186585

**Authors:** Tuan Pham, Julie S. Haas

## Abstract

As multimodal sensory information proceeds to the cortex, it is intercepted and processed by the nuclei of the thalamus. The main source of inhibition within thalamus is the reticular nucleus (TRN), which collects signals both from thalamocortical relay neurons and from thalamocortical feedback. Within the reticular nucleus, neurons are densely interconnected by connexin36-based gap junctions, known as electrical synapses. Electrical synapses have been shown to coordinate neuronal rhythms, including thalamocortical spindle rhythms, but their role in shaping or modulating transient activity is less understood. We constructed a four-cell model of thalamic relay and TRN neurons, and used it to investigate the impact of electrical synapses on closely timed inputs delivered to thalamic relay cells. We show that the electrical synapses of the TRN assist cortical discrimination of these inputs through effects of truncation, delay or inhibition of thalamic spike trains. We expect that these are principles whereby electrical synapses play similar roles in processing of transient activity in excitatory neurons across the brain.

## Introduction

It is well known that thalamocortical (TC) neurons relay sensory information to the cortex. For example, sensory information from rodent whiskers is projected from trigeminal nuclei to the ventroposteromedial (VPM) nuclei and posteromedial (POm) nuclei in the ventrobasal (VB) complex of the thalamus (Harris, 1987; Alloway, 2008). From VPM and POm, afferent connections relay information about whisking to the barrels of the primary somatosensory cortex (Sherman & Guillery, 1996). Within each of these nuclei, whisker inputs are encoded by varied latencies or spike rates (Sosnik *et al.*, 2001; Yu *et al.*, 2015).

During thalamocortical sensory relay, TC neuronal activity is regulated by a sheet of GABAergic neurons in the thalamic reticular nucleus (TRN) (Houser *et al.*, 1980). Most neurons within the ventrobasal complex (VB) receive monosynaptic GABAergic inputs from TRN neurons (Pinault & Deschenes, 1998), and there is strong monosynaptic excitation from VB to TRN (Ohara & Lieberman, 1985; Shosaku, 1986; Gentet & Ulrich, 2003). These reciprocal excitatory-inhibitory connections between TC and TRN neurons are likely to affect TC spiking, and the information relay from TC to cortex. For instance, large GABAergic conductances from TRN neurons have been shown to diminish information transfer in both computational models (Mayer *et al.*, 2006) and hybrid circuits of a model TRN – biological TC pair (Le Masson *et al.*, 2002).

Although TRN neurons are homogeneously GABAergic, the prevailing evidence suggests that intra-TRN inhibition is not prevalent in adult mice (Hou *et al.*, 2016). At the microcircuit level, GABAergic synapses are reported at less than 1% of nearby TRN pairs (Landisman *et al.*, 2002). The dominant source of intra-TRN connectivity, at least between nearby neurons, is thus electrical coupling via connexin36 (Cx36) gap junctions (Landisman *et al.*, 2002). Hence, sensory information relay to cortical neurons from TC neurons is regulated by both GABAergic feedback inhibition from TRN neurons and the electrical synapses between them.

Electrical synapses have been widely reported to participate in the generation of synchronous or phase-locked neuronal activity (Connors & Long, 2004). This role has been confirmed through models of networks with embedded electrical synapses (Lewis & Rinzel, 2003; Pfeuty *et al.*, 2005; Gutierrez *et al.*, 2013). Within thalamocortical circuits, electrical synapses of the TRN help to synchronize the spindle rhythms associated with slow-wave sleep or absence epilepsy (Lewis *et al.*, 2015; Fogerson & Huguenard, 2016). However, the role of electrical synapses in TRN on the transient, stimulus-evoked TC activity is relatively underexplored.

Here we examine the role of electrical synapses within TRN on the activity patterns of TC neurons. We use a reduced model of four cells: two pairs of reciprocally connected TC-TRN neurons, with an electrical synapse between the two TRN neurons. We deliver closely-timed inputs, mimicking inputs of similar temporal, spatial, or frequency arriving from sensory surround, to the TC cells. We examine how TC spiking is impacted by the inhibition delivered from the coupled TRN neurons. Our results demonstrate that electrical synapses can either fuse or further separate input-generate spiking, and we predict that these effects ultimately impacting the ability of recipient cortical cells to discriminate between the inputs.

## Methods

### 1. Model and simulation

Our model is based on a Hogkin-Huxley formulism for single compartmental model of TRN neurons (Destexhe *et al.*, 1996):

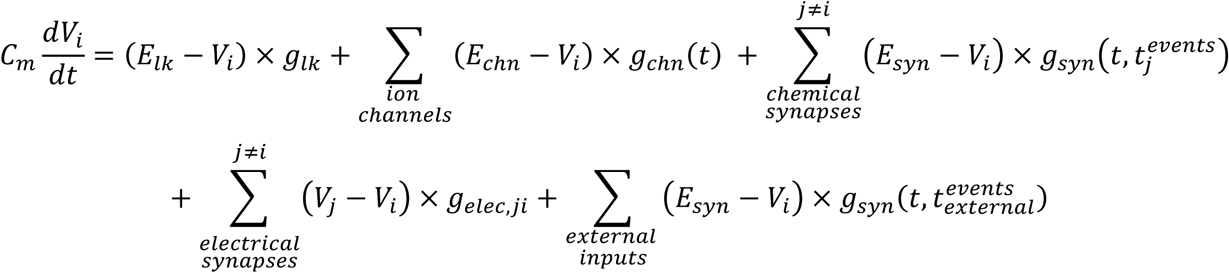

We used *C*_*m*_ of 1 μF/cm^2^. Ionic currents (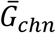, *E*_*chn*_) include fast transient Na^+^ current (60.5 mS/cm^2^, 50 mV); K^+^ delayed rectifier (60 mS/cm^2^, −100 mV); K^+^ transient A current (5 mS/cm^2^, −100 mV); slowly inactivating K-current K2 (0.5 mS/cm^2^, −100 mV); slow anomalous rectifier (H current) (0.025 mS/cm^2^, −40 mV); low threshold transient Ca^2+^ current (T current) (0.67 mS/cm^2^, 125 mV); leak current (0.06 mS/cm^2^, −75 mV). The membrane voltage initial condition (V0 = −70.6837 mV) was found by looking for the steady state after a simulation of 5000ms.

Chemical synapses include fast inhibitory GABA_A_ (*E*_*GABA*_ = −75 *mV*) and excitatory AMPA (*E*_*AMPA*_ = 0 *mV*) synapses. Synaptic conductance kinetics is implemented with a pair of fall and rise time constants (with *τ*_*rise*_ = 0.1 *τ*_*fall*_ for both synapses); *τ*_*fall,GABA*_ = 5*ms* and *τ*_*fall,AMPA*_ = 2*ms*. Implementation of synapses resembles NEURON’s implementation of (Traub *et al*, 2005) from ModelDB.

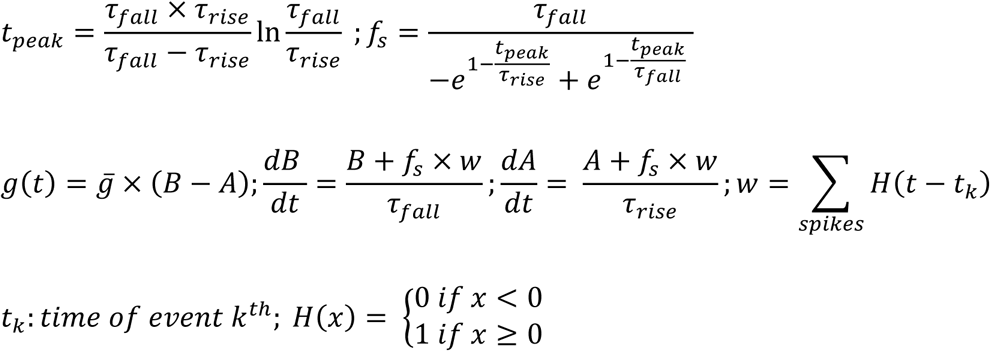

Our simulated network consists of 2 TRN cells connected via a single electrical synapse, and 2 TC cells receiving external input (Fig. 1A), simulated for 250ms. Within the network, TRN cells each send inhibitory input to TC cells via GABAA synapses, and TC cells send excitatory inputs to TRN cells via AMPA synapses. Since the model was easily excitable, no additional DC current was sent to TRN neurons to reach subthreshold excitation. External inputs are AMPAergic excitatory inputs and provided only to the two TC cells.

**Figure 1.**
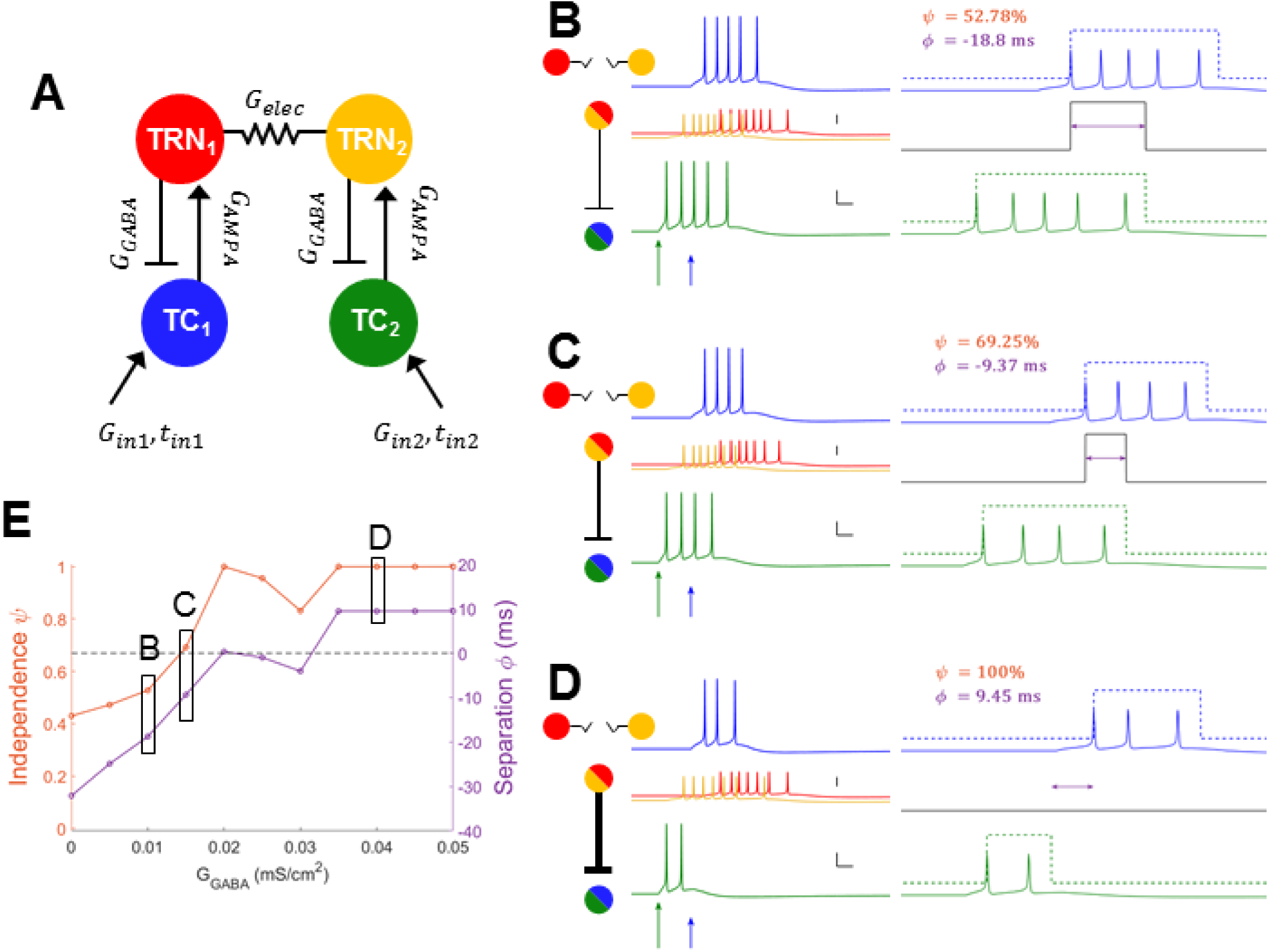
Model schematic and results with no electrical synapse. **A**: Model is composed of two sets of TRN and TC cells, with reciprocal chemical synaptic connections between pairs, and one electrical synapse between the two TRN cells. Each simulation of the model used different values of connection strengths G_elec_, G_GABA_ and input to TC_2_, G_in2_ and t_in2_. **B, C, D**: Simulation examples, each with G_elec_ = 0, and input to TC_2_ (green arrow) was large and early (G_in2_ = 0.09 mS/cm^2^, t_in2_ = 40 ms). Input to TC_1_ was constant (0.06 mS/cm^2^, 60 ms). Over the three simulations shown, inhibitory synapses increased in strength (G_GABA_ = 0.010, 0.015, 0.040 mS/cm^2^ respectively). In each subpanel, on the left are voltage traces color-coded as in (A), with the TRN traces superimposed and vertically reduced. Scale bars for TC traces are 20 mV, 10 ms; vertical scale bar of TRN traces is 40 mV. TC traces are expanded on the right, to demonstrate the computations for TC independence ψ and separation ϕ. Dotted lines represent the spiking window used to compute independence and separation. The black solid line indicates the amount of temporal overlap of TC spiking windows; purple arrows indicate the amount of temporal separation or overlap. **E**: Independence (orange) and separation (purple) shown for this input over all values of G_GABA_, with letters indicating the examples shown in B, C and D. The dashed black line marks the transition from overlapping spike trains (negative separation) and completely independent spike trains (positive separation).

Model electrical synapses (gap junctions) are linear and symmetrical. Values of electrical synapse conductance (*G*_*elec*_) varied from 0 to 0.025 mS/cm^2^, which converts to a coupling coefficient of roughly 0.2883. An applied current, iDC, was used from 0 to - 0.1μA/cm^2^ to quantify membrane conductance *G*_*m*_ ≈ 0.0551 mS/cm^2^ and the effective coupling coefficient values, which do not differ significantly with the theoretical values of *G*_*m*_ /(*G*_*m*_ + *G*_*elec*_) (data not shown).

The maximal GABAergic conductance (*G*_*GABA*_) varied from 0 to 0.05 mS/cm^2^, and the maximal AMPAergic conductance (*G*_*AMPA*_) between TC and TRN cells was fixed at 0.05 mS/cm^2^ to make sure that at least two spikes from a TC cell were required to elicit response from a TRN cell in an uncoupled network. External input to TC_1_ (*in*_1_) stays constant at maximal conductance (*G*_*in*1_) of 0.06 mS/cm^2^, and arrives (*t*_*in*1_) at 60ms. The arrival times (*t*_*in*2_) of external input to TC_2_ (*in*_2_) varied between 10 and 110 ms. The maximal conductance (*G*_*in*2_) of the input to TC_2_ varied from 0.02 to 0.1 mS/cm^2^.

Each simulation was simulated in parallel using MATLAB R2016 Parallel Toolbox and Lehigh University High Performance Computing Resources (Sol), solved with MATLAB’s *ode23* Runge-Kutta implementation using a maximum timestep of 0.01 ms.

### 2. Analysis

For each simulation, the number of spikes in each TC neuron was extracted. Temporal independence (*ψ*) and spiking separation (*ϕ*) were defined as outlined below.

Spiking window (*σ*) of a neuron was defined to be the temporal range between its first spike and last spike, with addition of a 5-ms window following the last spike used to allow for EPSP decay in a cortical cell.

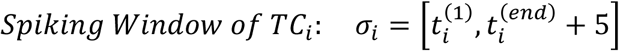

From each TC spike train, relative independence of the corresponding TC was calculated and normalized to values between 0 (complete overlap) and 1 (no overlap) as:

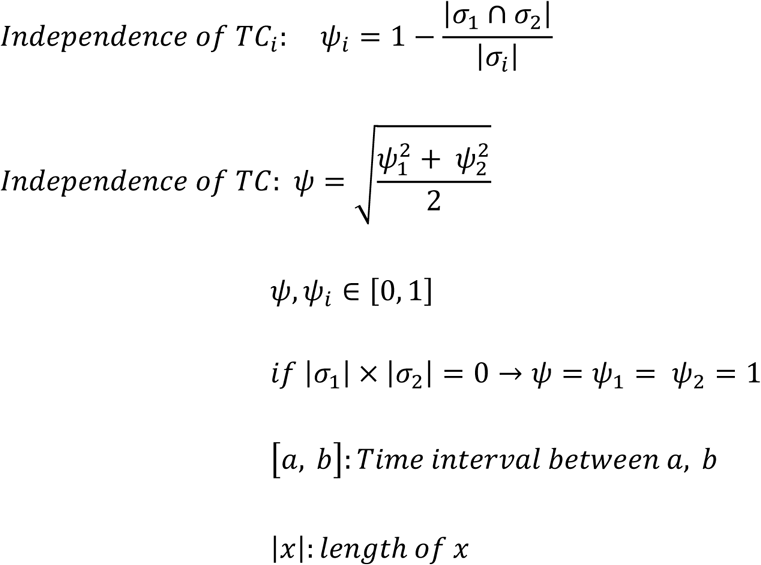

The separation of TC_1_ and TC_2_ spike trains was computed as the time difference between the spiking window termination of the leading neuron and the spiking window beginning of the following neuron. In cases of either TC_1_ or TC_2_ not spiking, this measure was not calculated. In overlapping cases, this measure is artificially considered to be negative, as loss of separation, and the magnitude is the amount of overlap.

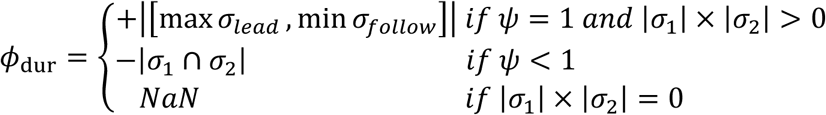

## Results

To examine the impact of electrical synapses of the TRN on thalamocortical transmission, we constructed a 4-cell model comprising two pairs of reciprocally connected TC and TRN cells, with an electrical synapse between the two TRN cells (Fig. 1A). Input delivered to TC_1_ was fixed at a constant arrival time and strength, and the inputs to TC_2_ were varied in arrival time and input strength. We used input strengths that resulted in a burst of spikes that is typical of thalamic cells, lasting for tens of ms (e.g. Fig. 1B). To characterize the results of electrical synapses on the output of the TC cells, we quantified independence (ψ) as the percentage as inversely related to the temporal intersection of the two TC spike trains, normalized to the duration of spiking. We also quantified separation (ϕ) as the time interval between the termination of one spike train and the onset of spiking in the other TC cell; negative values of separation result from overlapping trains. Higher values of independence and positive values of separation, we believe, increase the chances that a cortical cell that receives inputs from both of these TC cells will be able to discriminate the input as arising from different pathways (e.g. whiskers).

With no electrical synapse, the inhibitory feedback from TRN to TC cells is the sole influence that acts to separate the spike trains and thus the inputs that TC relays to cortex (Fig 1B). More specifically, increases in inhibitory strength between TRN and TC cells result in earlier termination of spiking in the TC cells (Fig 1B-D), resulting in both increases in separation and independence of the trains from each other (Fig. 1E).

For increases in strength of the electrical synapse between TRN cells, we observed several effects that ultimately impacted the separation and independence of TC spiking: latency of spiking in the TC that received inputs later, truncation of spike trains, and prevention of any spiking. Electrical synapse strengths were matched to those observed in vitro for TRN cells (Landisman *et al.*, 2002; Haas *et al.*, 2011), with a maximal coupling coefficient ~0.3.

An example of GJ-mediated changes in latency is shown in Figure 2. In this case, input was delivered 40 ms to TC_1_ before the input in TC_2_, and both inputs were of the same strength. Inhibitory strength between TRN and TC cells was relatively low. While the spike trains are always independent in this case, the initial separation (without electrical synapses) is 9.2 ms (Fig. 2A_i_), and a cortical neuron that received inputs from both TC_1_ and TC_2_ may not differentiate between one spike train that starts 9.2 ms after the termination of the previous train. Increases in electrical strength within the model systematically resulted in delayed spiking in TC_2_ (Fig. 2A_i-iv_) with corresponding increases in separation, up to 30 ms (Fig. 2A_*iv*_), and ultimately prevented spiking in TC_2_ (Fig. 2A_*v*_). In these cases, separation values follow latencies, and show a strong dependence on both electrical and inhibitory synapse strengths (Fig. 2B, 2D). Electrical synapse strength acts synergistically with inhibitory synapse strength (Fig. 2E), such that increases in both parameters result in large latency changes, or ultimately prevention of spiking in TC_2_ (whitespace in Fig. 2D and 2E).

**Figure 2.**
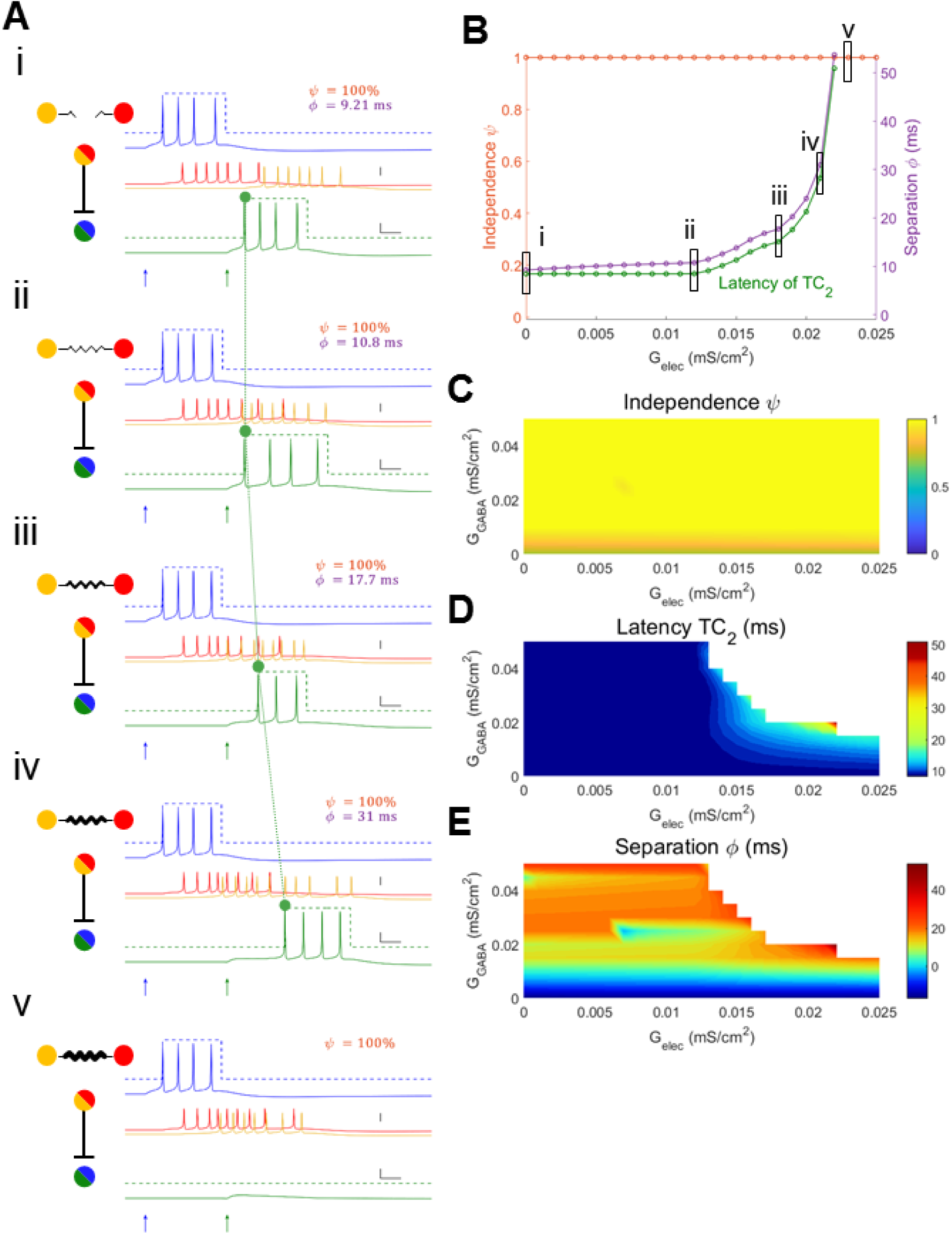
Electrical synapses between TRN cells result increase latency of TC spiking. **A:** Example simulations with superimposed TC spiking windows. Input to TC_1_ was constant (0.06 mS/cm^2^, 60 ms). For these cases, input to TC_2_ was 0.06 mS/cm^2^, 100 ms, and inhibitory synapses were G_GABA_ = 0.020 mS/cm^2^. Subpanels *i* – *v* show simulations with increasing electrical synapse strengths (G_elec_ = 0, 0.012, 0.018, 0.021, 0.023 mS/cm^2^ respectively); for v, separation is undefined because at least TC_2_ does not spike. **B**: Independence (orange), separation (purple) and TC_2_ latency (green) for simulations in A, over all values of electrical synapse strength. Latency of TC_2_ has the same axis as separation. **C, D, E**: Heat maps for independence, TC_2_ latency, separation respectively against G_elec_ and G_GABA_. Whitespace in latency indicates no spiking in TC_2_ while blank space in separation means there is no spiking in either TC cell.

Spike train truncation arises for stronger values of inhibition within the network, and is modulated by the electrical synapse (Fig. 3). As electrical synapse strength increases, spiking rate in TC_2_ decreases (Fig. 3A). Independence of the TC spike trains varies with increases in electrical synapse strength, as the spike train of TC_2_ diminishes (Fig. 3D), and separation also increases (Fig. 3E). In this case, the baseline condition (with no or weak electrical synapses) is moderately overlapping. However, the strong inhibition in this set acts through increasing electrical synapse strengths to prevent prolonged spiking of the first spike train (Fig 3A_i-iii_), effectively decreasing overlap, thereby increasing both independence and separation. For varied values of inhibitory strength within the network, the relationships between independence, separation and electrical synapses become more complex, resulting from the dependence of rate (Fig. 3D) on both electrical and inhibitory synapse strength.

**Figure 3.**
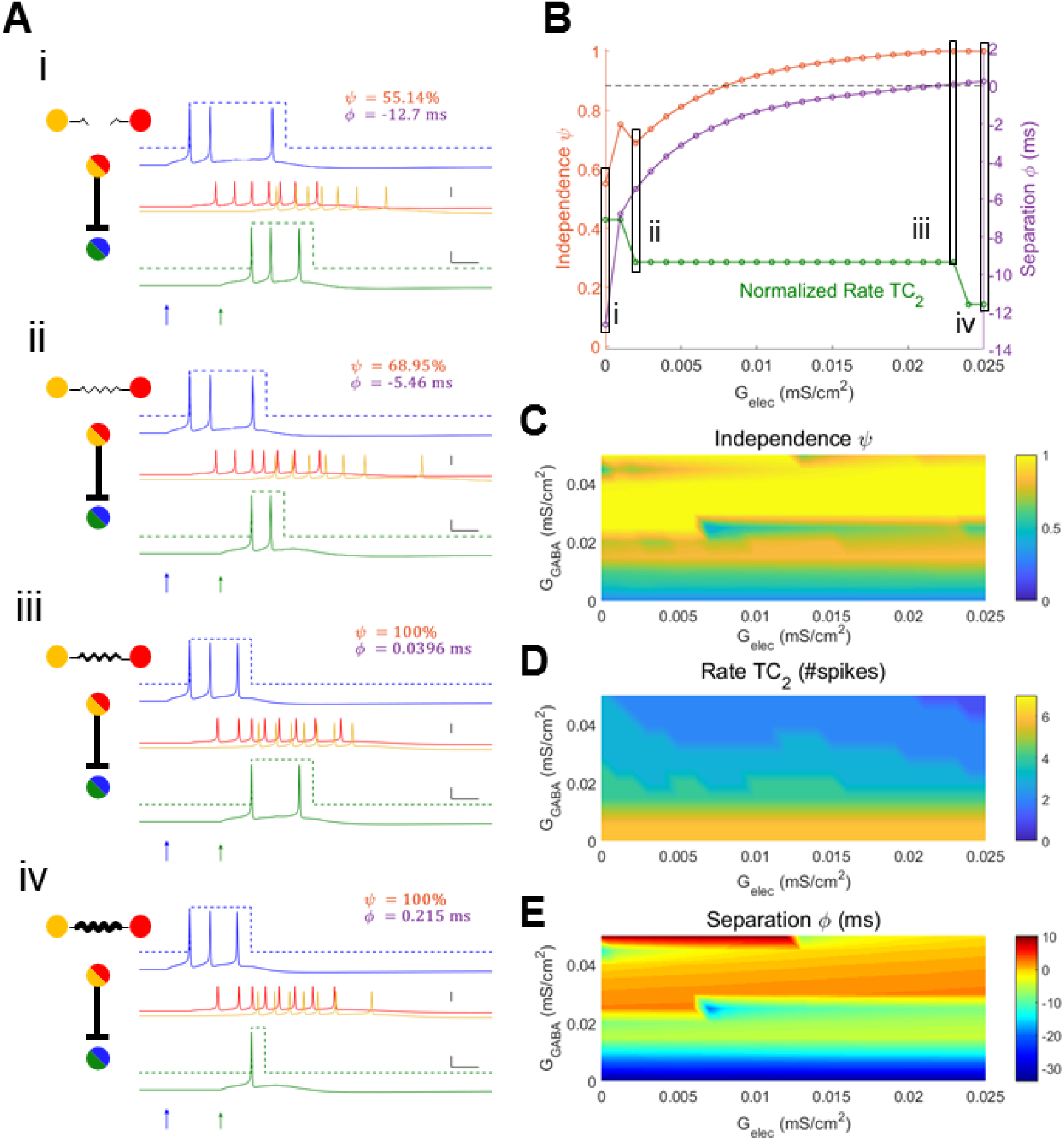
Electrical synapses between TRN cells result in truncated TC spike trains. **A**: Example simulated traces. Input to TC_1_ was constant (0.06 mS/cm^2^, 60 ms). For these cases, input to TC_2_ was 0.05 mS/cm^2^, 80 ms and inhibitory synapses were G_GABA_ = 0.045 mS/cm^2^; Subpanels *i* – *iv* show simulations with increasing electrical synapse strengths (G_elec_ = 0, 0.002, 0.023, 0.025 mS/cm^2^ respectively). **B**: Independence (orange), separation (purple) and normalized TC_2_ rate (green) for the set of simulations in (A) plotted against electrical synapse strength. Normalized rate of TC_2_ is the number of TC_2_ spikes relative to the maximum over all simulations (7 spikes). **C, D, E**: Heat map for independence, TC_2_ rate (unnormalized), and separation respectively plotted against G_elec_ and G_GABA_.

In general, spiking rate depends on both inhibitory and electrical synapse strength, as well as on the details of the input, specifically on its arrival time and strength. The dependence of spiking rate in TC_2_ on electrical and inhibitory synapse strength is shown in Figure 4. Generally, electrical synapses between TRN cells are more effective in terminating trains for more temporally separated inputs (Fig. 4, top row), as the delays for input integration in TRN cells and TC cells determine the timescale at which electrical synapses influence the circuit. The more different the inputs are, either in time or amplitude, the larger the effects are from the electrical or the inhibitory synapse.

**Figure 4.**
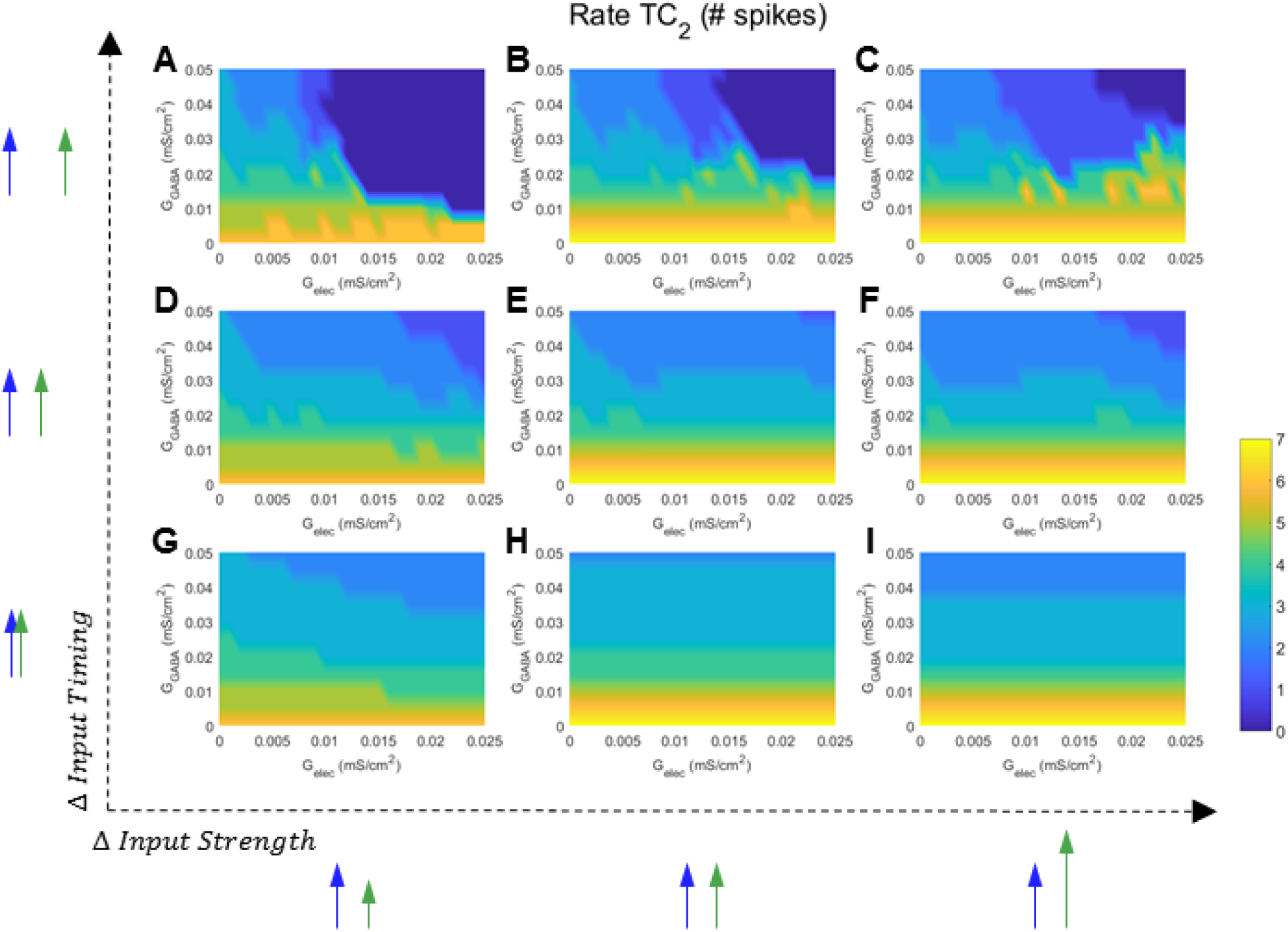
Electrical synapses between TRN cells modulate rate of TC_2_. Each panel (**A – I**) is a heat map for TC_2_ rate, plotted against all values of G_elec_ and G_GABA_. Input strengths to TC_2_ were varied between panels from left to right (G_in2_ = 0.04, 0.06 and 0.08 mS/cm^2^; input to TC_1_ was always G_in1_ = 0.06 mS/cm^2^), and input timing was varied from bottom to top (t_in2_ = 60, 80 and 100 ms; t_in1_ was always 60 ms).

Electrical and inhibitory synapses also act together to regulate spike train separation (Fig. 5). As for spiking rate, the biggest impact of electrical synapses on separation is seen for inputs that are different in arrival time (Fig. 5A-C).

**Figure 5.**
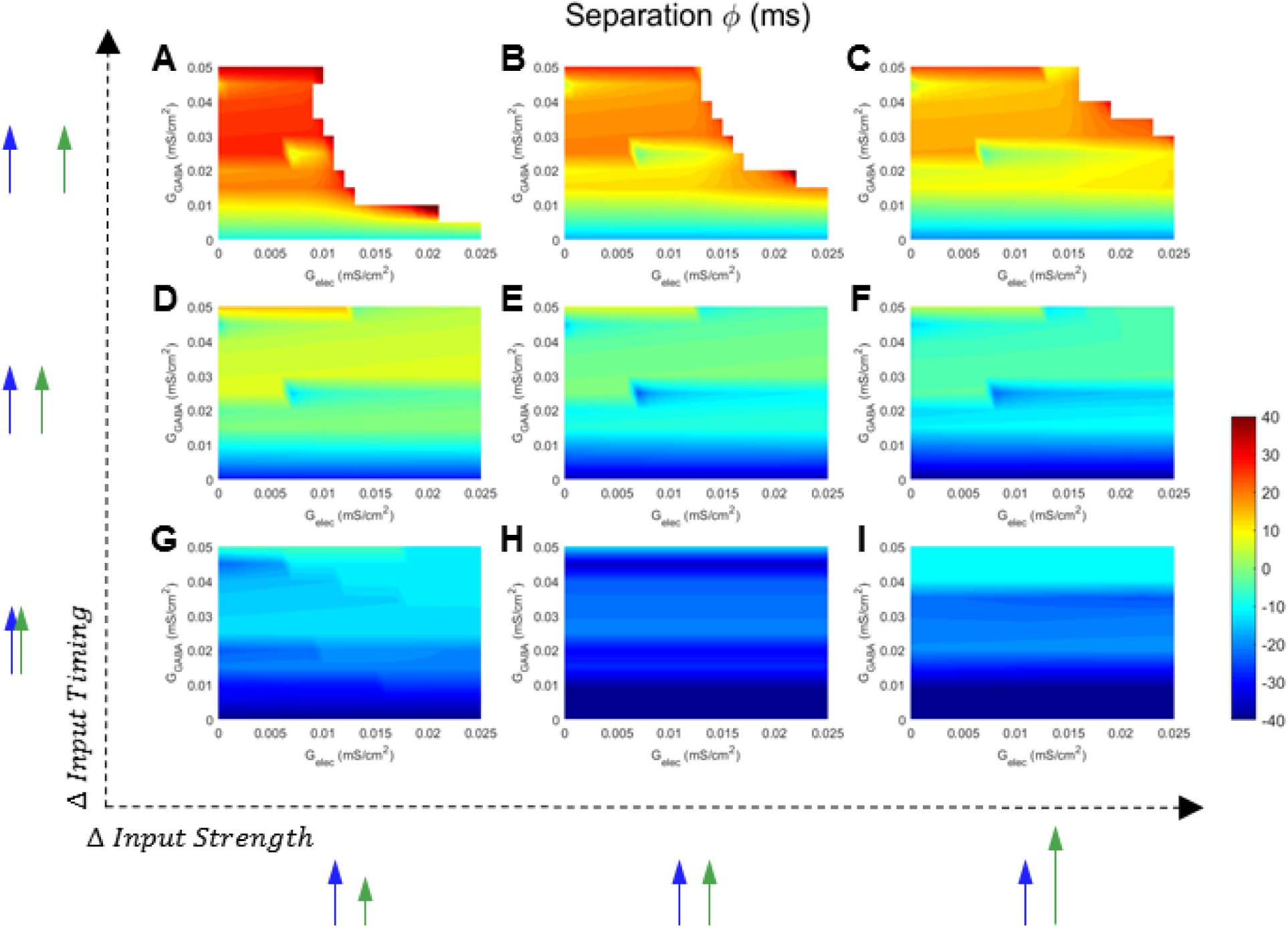
Electrical synapses between TRN cells increase separation of TC spike trains. Each panel (**A – I**) is a heat map for separation between TC spike trains, plotted against all values of G_elec_ and G_GABA_. Input strengths to TC_2_ were varied between panels from left to right (G_in2_ = 0.04, 0.06 and 0.08 mS/cm^2^; input to TC_1_ was always G_in1_ = 0.06 mS/cm^2^), and input timing was varied from bottom to top (t_in2_ = 60, 80 and 100 ms; t_in1_ was always 60 ms). Whitespace in separation means there is no spiking in either TC cell.

For closely timed and similar inputs, electrical synapses (or inhibitory synapses) are relatively ineffective in separating the inputs. In fact, for similar inputs, we observed that electrical synapses, acting through inhibitory synapses, instead increase temporal fusion of the inputs. This effect is shown in the progressive decrease in independence, indicated by the increase in the area of blue shading across sets of panels, in Figure 6. Increases in electrical synapse strength, in this context, broaden the set of input differences for which spike trains are non-independent, seen when comparing panels within Fig. 6B.

**Figure 6.**
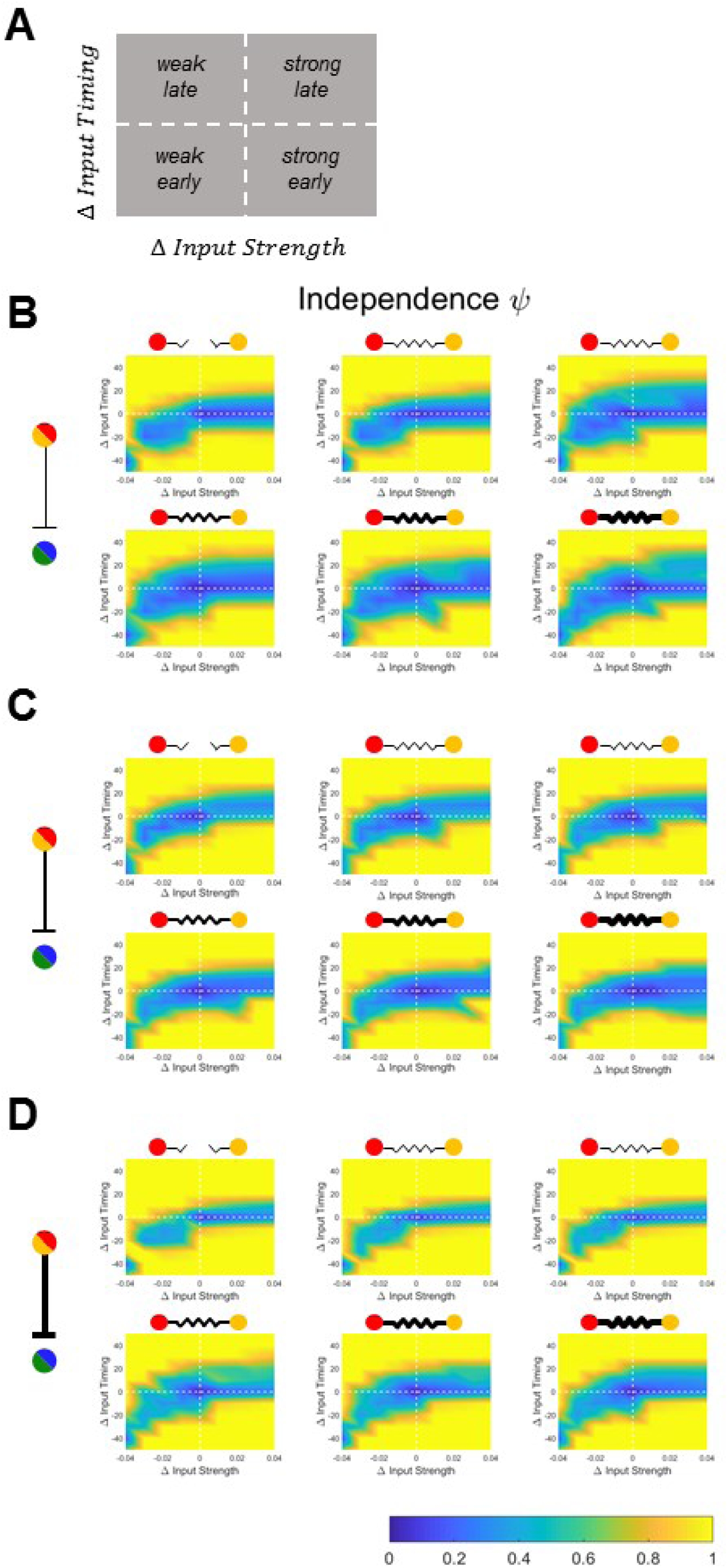
Electrical synapses between TRN cells merge TC spike trains for inputs that are similar in timing and strength. **A**: Display convention for the following results, plotted by the differences of inputs to TC_2_ in arrival time and strength relative to the fixed input to TC_1_. The horizontal and vertical dashed lines represent simultaneous and equal inputs, respectively. **B, C, D**: Independence plotted for varied values of electrical synapse strength and GABAergic inhibition (G_GABA_ = 0.025 in **B**, 0.040 in **C**, 0.050 mS/cm^2^ in **D**). The electrical synapse strength is indicated by thickness of symbol (G_elec_ = 0, 0.005, 0.010, 0.015, 0.020, 0.025 mS/cm^2^).

Finally, the net differences in independence created by electrical synapses are shown in Fig. 7. In this plot, each subpanel is shown as the difference in spike train independence between a set of simulations, and the baseline set of simulations when electrical synapses are absent. As in Fig. 6, identical inputs are the center of each subpanel. From this figure, we see that as electrical synapse strength is increased (left to right), the gains in independence from baseline of spike trains in the two TC cells can both increase or decrease (yellow or blue). Together, our results show that electrical synapses between TRN neurons add a variety of possible outcomes to TC spike trains, compared to simple feedback inhibition.

**Figure 7.**
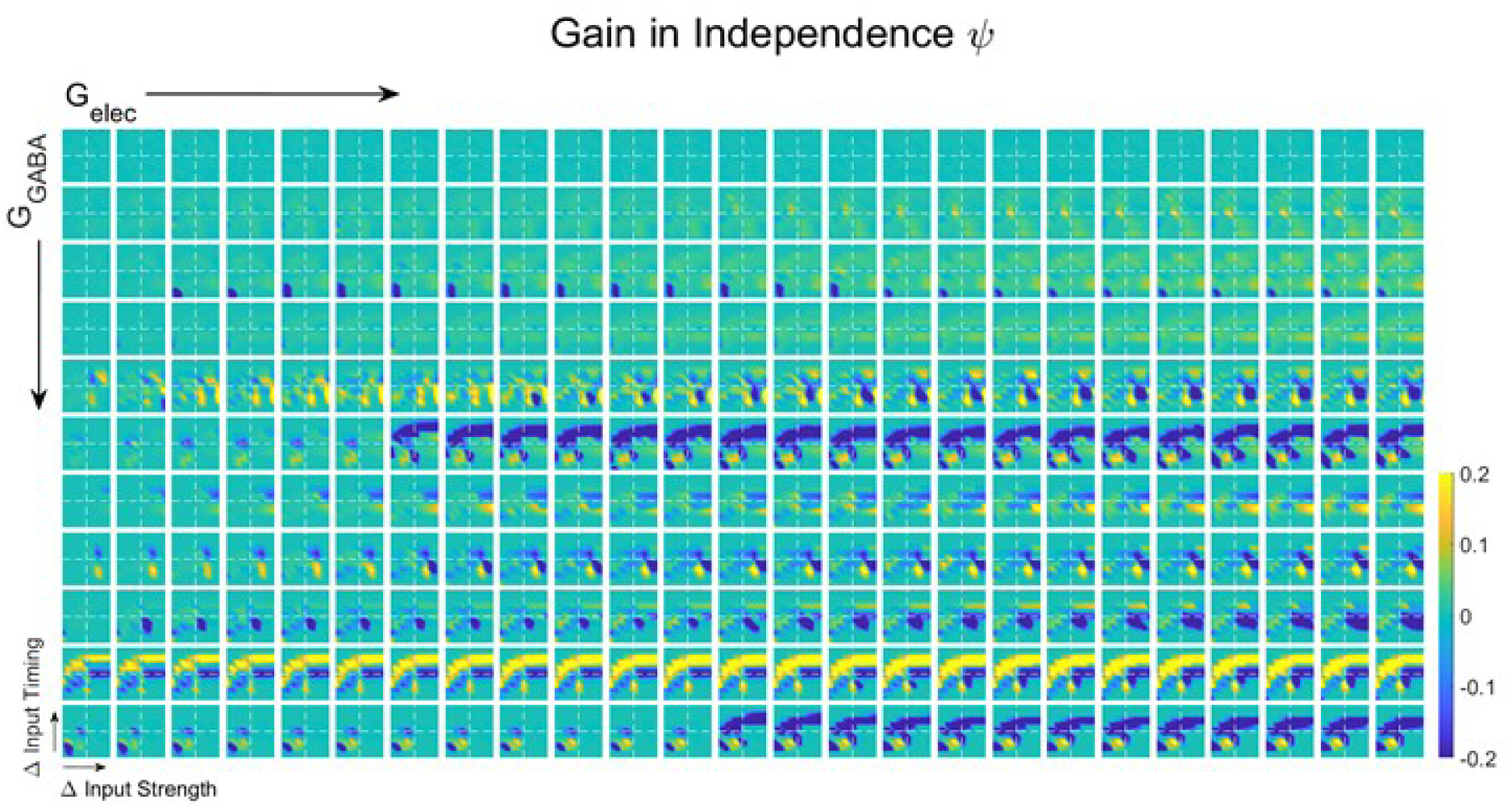
Change in TC spiking independence, compared to unconnected (G_elec_ = 0) baseline, resulting from electrical synapses between TRN cells. Each subpanel is the difference of TC spike train independence, between the presence of electrical synapse (G_elec_ ≠ 0) and the unconnected case (G_elec_ = 0) with similar inhibitory conductance. Within each subpanel, input strength and size are represented as in Figure 6, with horizontal and vertical dashed lines representing simultaneous and equal inputs, respectively. From left to right, electrical synapse conductance increases across panels (G_elec_ = 0.001 to 0.025 mS/cm^2^). From top to bottom, inhibitory synapse conductance increases (G_GABA_ = 0 to 0.050 mS/cm^2^).

## Discussion

Noting that most of the experimental demonstrations or computational simulations of electrical synapses focus on the relationship between electrical synapses and synchrony, we set out to explore the impact of electrical synapses on transient spike train processing within the brain. Using a minimal 4-cell model of paired thalamus and thalamic reticular nucleus cells with a single electrical synapse to connect the pairs, here we have shown that through feedback inhibition, electrical synapses of the TRN modify both timing and rate of thalamic spiking. Ultimately, the electrical synapses of the TRN modulate both the temporal independence and separation of the spike trains that thalamic cells send on to cortex, thus impacting whether cortical cells receive spike trains that can be discriminated as arising from separate receptors or receptive fields of the sensory surround.

The results of our simulations are important in the context of plasticity of electrical synapses in the TRN (Landisman & Connors, 2005; Haas *et al.*, 2011). Our results show that the strength of electrical synapses can shift the character of a relay network, from one that separates its inputs to one that fuses inputs, for instance. Changes in the strength of electrical synapses are represented here by a shift along an axis, and show that even smaller changes in electrical synapse strength have the potential to change output rate, timing, independence and separation.

Our model is the simplest core unit, or motif, of intrathalamic connectivity. Previous results have shown that the reciprocal connections within a single TRN-TC pair can strongly depress the process of thalamocortical sensory relay (Le Masson *et al.*, 2002). Our intrathalamic motif with two coupled pairs further shows that electrical synapse not only affects thalamocortical rates and latencies, thus contributing to basic coding of spatial-temporal sensory inputs (for example, whisker inputs (Sosnik *et al.*, 2001; Yu *et al.*, 2015)), but also regulates temporal independence and separation. Changes in separation due to electrical synapses might also increase efficient coding, by modulating the sparsity of sensory inputs and relay.

Our model is the simplest core unit, or motif, of intrathalamic connectivity, and our model assumed these four cells to be identical in term of intrinsic properties. However, connections within thalamus and between thalamus and cortex are more complex. Thalamic cells receive convergent inhibitory inputs from multiple TRN cells, and TRN cells receive input from multiple thalamic relay cells. While we have focused on straightforward thalamic relay of singular inputs, mimicking POm, other thalamic subsectors receive inputs from broad areas of the sensory surround. Cortical feedback is also not represented in the present model. Thus, there is much future work to be done to thoroughly explore the role of electrical synapses in transient signal processing within thalamus.

The underlying circuit – inhibitory neurons connected by an electrical synapse, and providing feedback inhibition to the principal neurons that excite them – is one that we expect may be embedded within retina, where AII amacrine cells regulate retinal ganglion cell spiking, and within cortex, where inhibitory neurons regulate principal cell firing. We expect that the general principles seen here – that electrical synapses act through inhibitory synapses to increase latency, decrease spike rates, and modify the independence and separation of spike trains in principal cells – will also impact information processing in the many areas that contain electrical synapses.

